# Depurination of colibactin-derived interstrand cross-links

**DOI:** 10.1101/869313

**Authors:** Mengzhao Xue, Kevin M. Wernke, Seth B. Herzon

## Abstract

Colibactin is a genotoxic gut microbiome metabolite long suspected of playing an etiological role in colorectal cancer progression. Evidence suggests colibactin forms DNA interstrand cross-links (ICLs) in eukaryotic cells and activates ICL repair pathways, leading to the production of ICL-dependent DNA double-strand breaks (DSBs). Here we show that colibactin ICLs can evolve directly to DNA DSBs. Using the topology of supercoiled plasmid DNA as a proxy for alkylation adduct stability, we show that colibactin-derived ICLs are unstable toward depurination and elimination of the 3′ phosphate. This pathway leads progressively to the formation of nicks SSBs and cleavage DSBs and is consistent with the earlier determination that non-homologous end joining repair-deficient cells are sensitized to colibactin-producing bacteria. The results herein further our understanding of colibactin-derived DNA damage and underscore the complexities underlying the DSB phenotype.

## Introduction

Colibactin is a genotoxic microbiome gut metabolite that has been implicated in the pathogenesis of colorectal cancer.^1^ The biosynthesis of colibactin is encoded in a 54-kb biosynthetic gene cluster (BGC) referred to as *clb* or *pks* and is widely distributed among *Enterobacteriaceae*.^2^ The *clb* gene cluster has undergone heavy scrutiny by the microbiome community due to its suspected role in human colorectal cancer (CRC) progression. *E. coli* possessing the *clb* gene cluster are found in 55–67% CRC patients^3,4^ and accelerate progression from dysplasia to invasive carcinoma in mouse models of intestinal inflammation.^3^ This evidence suggests that colibactin, the metabolite produced by *clb*, is the agent underlying tumorigenesis.^1^ The structure of colibactin was recently proposed as **1** by isotope labeling and tandem MS, and confirmed by chemical synthesis (Fig. 1).^5^ Colibactin (**1**) contains two cyclopropane residues appended to an α,β-unsaturated imine (blue in **1**). The unsaturated imines activate the cyclopropane rings, allowing for alkylation of DNA by nucleotide addition (see **1**→**2**, Fig. 1).^6^

**Fig. 1.**
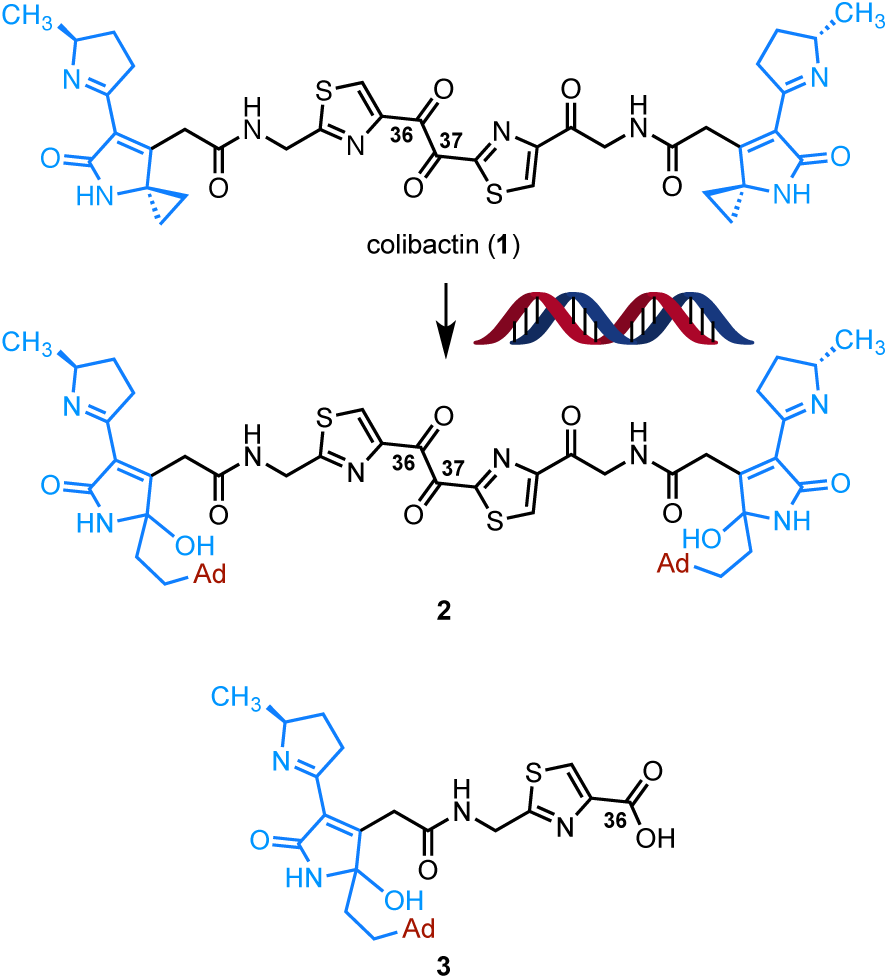
Structure of colibactin (**1**), the diadenine adduct **2**, and the first detected monoadenine adduct **3**.

*Clb*^+^ *E. coli* are genotoxic. In 2006 Oswald and co-workers established that HeLa cells infected with *clb*^+^ *E. coli* accumulated DNA double-strand breaks (DSBs) and lagged in S phase.^7^ More recently, these researchers showed that DNA interstrand cross-links (ICLs) appear to be the dominant cellular phenotype of *clb*^+^ *E. coli*.^8^ Thus, *clb*^+^ *E. coli* were found to form ICLs in linearized plasmid DNA or cellular genomic DNA, and to activate ICL repair pathways. Consistent with this, the bis(adenine) adduct **2** was characterized using mass spectrometry and isotope labeling when linearized plasmid DNA was treated with synthetic colibactin (**1**)^5^ or *clb*^+^ *E. coli.*^5,9^ The ICL and DSB phenotypes are interrelated. ICLs are resolved by the Fanconi Anemia (FA) pathway during S phase^10^ and repair is initiated by convergence of two replication forks on either side of the ICL. In subsequent steps, the ICL is excised from one parent strand resulting in a DNA DSB. This DSB is resolved by homologous recombination (HR) repair, one of the two primary DSB repair pathways.^11^ Thus, HR repair markers, such as phospho-SER139-H2AX (γH2AX), are observable when DSBs are formed during FA repair.^12^

However, in their original report Oswald and co-workers noted that cells deficient in non-homologous end joining (NHEJ), the second major DSB repair pathway (which operates throughout the cell cycle), were sensitized to *clb*^+^ *E. coli*.^7^ Because FA is coupled to HR but not NHEJ, this suggests generation of FA-independent DSBs. In our studies of synthetic colibactin mimetics^6^ and presently colibactin (**1**) itself, we observed fragmentation of alkylated DNA during denaturing electrophoresis. The mononadenine adduct **3** has been characterized by NMR spectroscopy and was shown as linked through N3 of adenine.^13^ N3-Adenine alkylation products are susceptible to depurination,^14^ and the resulting apurinic (AP) sites can progress to SSBs by elimination of the 3′ phosphate (shown in Fig. 6 below).^15^ Alternatively, AP sites are recognized by apurinic/apyrimidinic endonuclease 1 (APE1), which cleaves the phosphate 5′ to an abasic site, leaving a free 3′ hydroxyl and 5′ deoxyribose phosphate. We hypothesized that 3′ phosphate elimination and/or APE1 activity may contribute to the formation of DSBs outside of S phase. To test this hypothesis, we performed experiments to probe the stability of ICLs generated by *clb*^+^ *E. coli* and synthetic colibactin. Our data suggest DSBs derived from colibactin may arise from a combination of FA repair and depurination of ICLs.

**Fig. 2.**
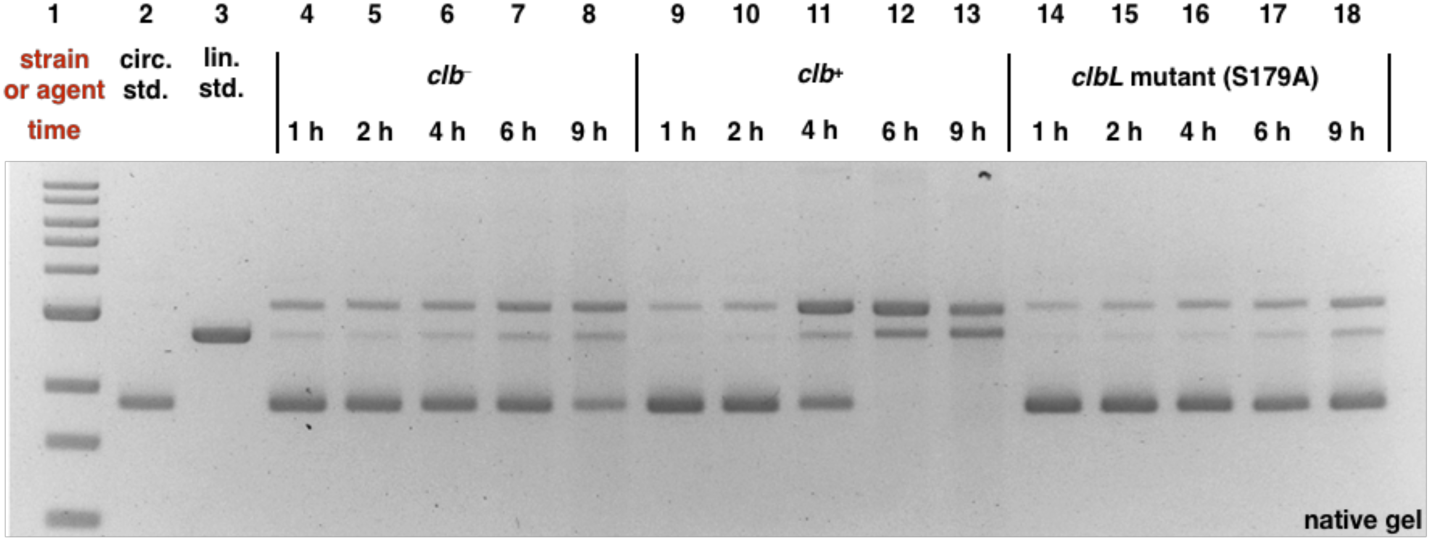
DNA plasmid cleavage assay employing circular pUC19 DNA and *E. coli* and analysis by neutral gel electrophoresis. DNA ladder (Lane #1); circular pUC19 DNA standard (Lane #2); linearized pUC19 DNA standard (Lane # 3); *clb*^−^ BW25113 *E. coli* 1 h (Lane #4), 2 h (Lane #5), 4 h (Lane #6), 6 h (Lane #7), and 9 h (Lane #8); *clb*^+^ BW25113 *E. coli* 1 h (Lane #9), 2 h (Lane #10), 4 h (Lane #11), 6 h (Lane #12), and 9 h (Lane #13); *clbL* mutant (S179A) BW25113 *E. coli* 1 h (Lane #14), 2 h (Lane #15), 4 h (Lane #16), 6 h (Lane #17), and 9 h (Lane #18). Conditions (Lane #4–#18): *clb*^−^ BW25113 *E. coli* (Lane #4–#8), *clb*^+^ BW25113 *E. coli* (Lane #9–#13), and *clbL* mutant (S179A) BW25113 *E. coli* (Lane #14–#18), circular pUC19 DNA (7.7 µM in base pairs), M9-CA media, 37 °C, 1 h, 2 h, 4 h, 6 h, or 9 h. The DNA was isolated, purified, and analyzed by native agarose gel electrophoresis (90 V, 2 h). Order of elution (bottom to top): supercoiled plasmid DNA, linearized DNA, nicked DNA.

**Fig. 3.**
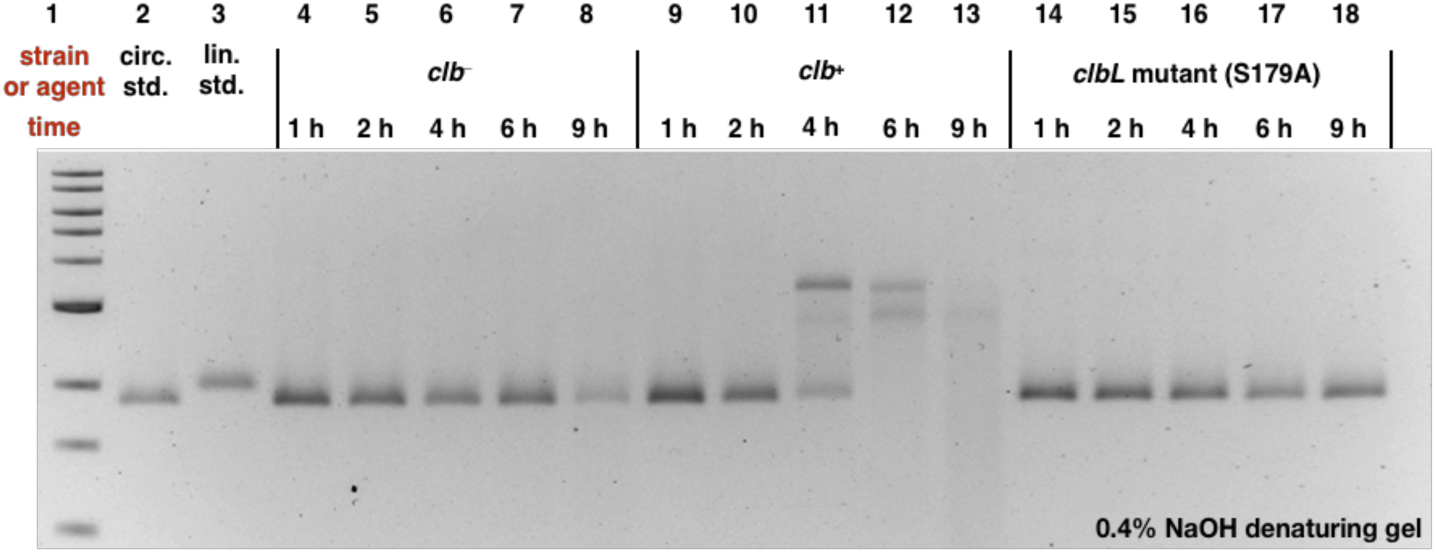
DNA plasmid cleavage assay employing circular pUC19 DNA and *E. coli* and analysis by denaturing electrophoresis. DNA ladder (Lane #1); circular pUC19 DNA standard (Lane #2); linearized pUC19 DNA standard (Lane # 3); *clb*^−^ BW25113 *E. coli* 1 h (Lane #4), 2 h (Lane #5), 4 h (Lane #6), 6 h (Lane #7), and 9 h (Lane #8); *clb*^+^ BW25113 *E. coli* 1 h (Lane #9), 2 h (Lane #10), 4 h (Lane #11), 6 h (Lane #12), and 9 h (Lane #13); *clbL* mutant (S179A) BW25113 *E. coli* 1 h (Lane #14), 2 h (Lane #15), 4 h (Lane #16), 6 h (Lane #17), and 9 h (Lane #18). Conditions (Lane #4–#18): *clb*^−^ BW25113 *E. coli* (Lane #4–#8), *clb*^+^ BW25113 *E. coli* (Lane #9–#13), and *clbL* mutant (S179A) BW25113 *E. coli* (Lane #14–#18), circular pUC19 DNA (7.7 µM in base pairs), M9-CA media, 37 °C, reaction proceed for 1 h, 2 h, 4 h, 6 h, and 9 h. The DNA was isolated, purified, and analyzed by 0.4% NaOH denaturing agarose gel electrophoresis (90 V, 2 h).

**Fig. 4.**
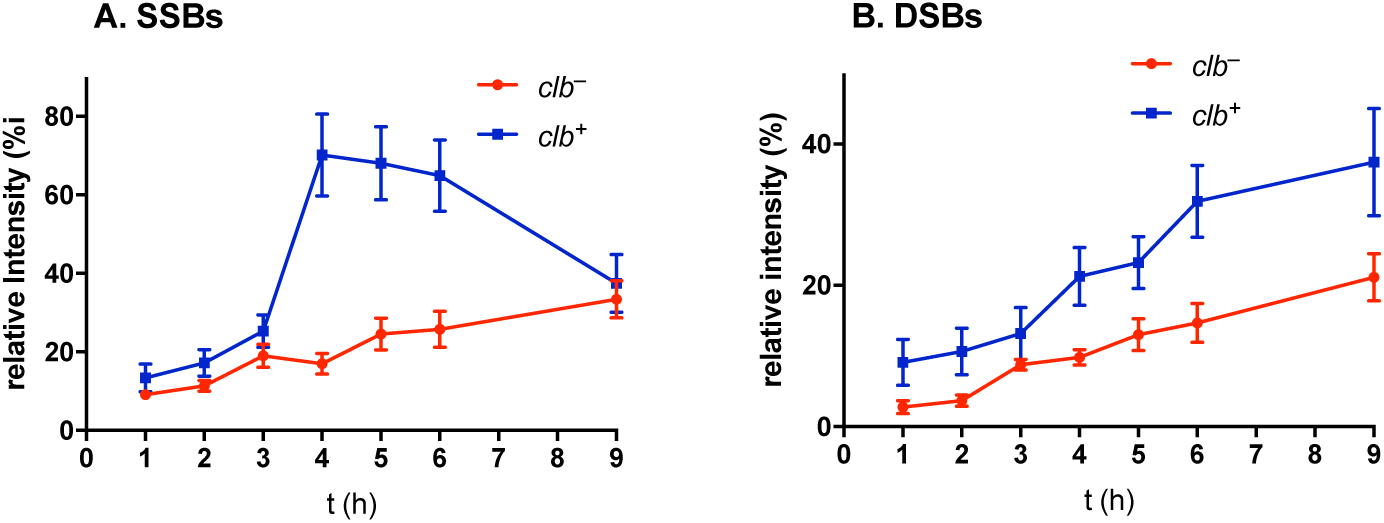
Formation of DNA SSBs and DSBs in circular pUC19 plasmid DNA treated with *clb*^+^ or *clb*^−^ *E. coli*. **A.** SSBs as a function of time. **B.** DSBs as a function of time. SSBs and DSBs are expressed as intensity of their respective bands relative to an internal control (untreated DNA). Three technical replicates (Fig. S2).

**Fig. 5.**
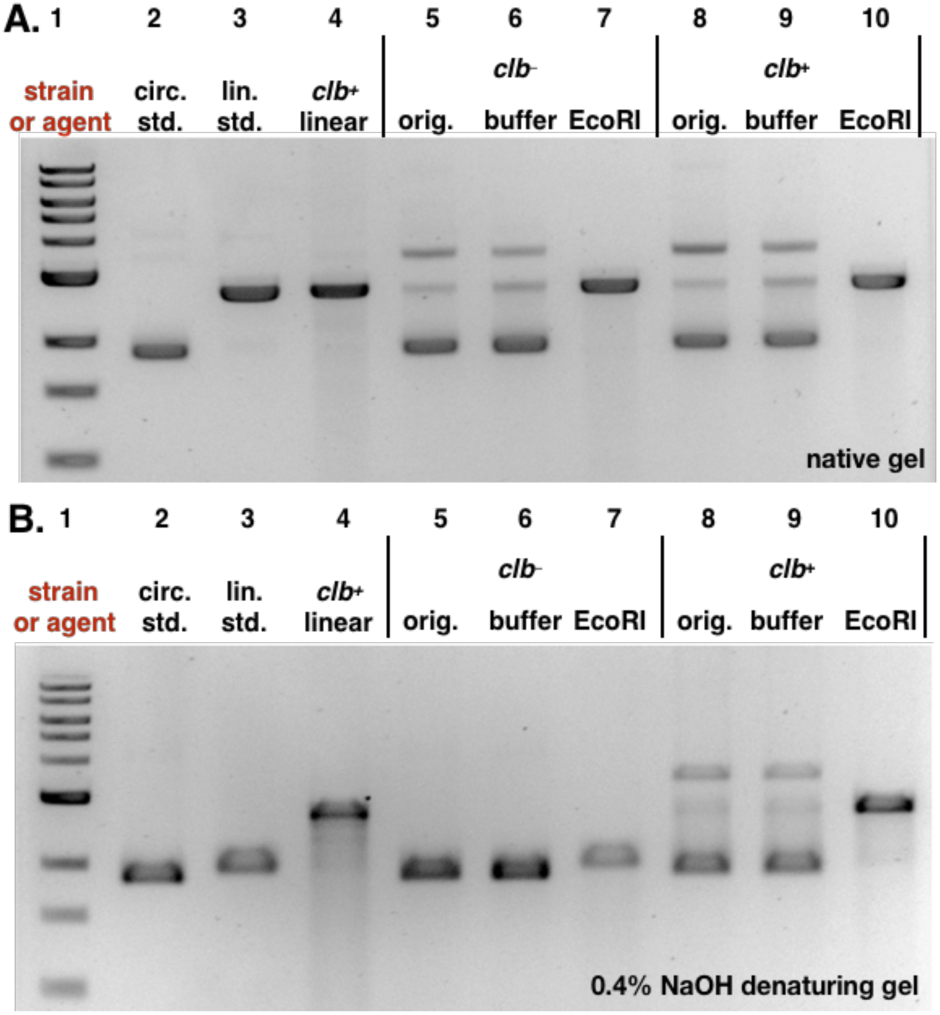
Analysis of pUC19 DNA following treatment with *clb*^−^ or *clb*^+^ *E. coli* and linearization with the restriction enzyme EcoRI. The cross-linked linearized pUC19 DNA isolated from a co-culture with *clb*^+^ BW25113 *E. coli* was used a positive control. **A.** Analysis of DNA by native gel electrophoresis. **B.** Analysis of DNA by denaturing gel electrophoresis. For both A and B: DNA ladder (Lane #1); circular pUC19 DNA standard (Lane #2); linearized pUC19 DNA standard (Lane # 3); linearized pUC19 DNA co-cultured with *clb*^+^ BW25113 *E. coli* (Lane #4); circular pUC19 DNA isolated from co-culture with *clb*^−^ BW25113 *E. coli* (Lane #5), reacted with buffer (Lane #6), reacted with EcoRI restriction enzyme (Lane #7); circular pUC19 DNA isolated from co-culture with *clb*^+^ BW25113 *E. coli* (Lane #8), reacted with buffer (Lane #9), reacted with EcoRI restriction enzyme (Lane #10). Conditions (Lane #4): linearized pUC19 DNA, *clb*^+^ BW25113 *E. coli*, M9-CA media, 4 h at 37 °C. Conditions (Lane #5–#7): circular pUC19 DNA isolated from co-culture with *clb*^−^ BW25113 *E. coli* in M9-CA media for 4 h at 37 °C (Lane #5); the DNA (15.4 µM base pair) was reacted with CutSmart Buffer® (New England Biolabs®), pH 7.9, at 37 °C for 30 minutes (Lane #6); the DNA (15.4 µM base pair) was reacted with 20 units of EcoRI-HF restriction enzyme in CutSmart Buffer® (New England Biolabs®), pH 7.9, at 37 °C for 30 minutes (Lane #7). Conditions (Lane #8–#10): circular pUC19 DNA isolated from co-culture with BW25113 *clb*^+^ *E. coli.* in in M9-CA media for 4 h at 37 °C (Lane # 8); the DNA (15.4 µM base pair) was reacted with CutSmart Buffer® (New England Biolabs®), pH 7.9, at 37 °C for 30 minutes (Lane #9); the DNA (15.4 µM base pair) was reacted with 20 units of EcoRI-HF restriction enzyme in CutSmart Buffer® (New England Biolabs®), pH 7.9, at 37 °C for 30 minutes (Lane #10). The DNA was isolated and analyzed by native (Fig. 5A) or 0.4% NaOH denaturing (Fig. 5B) agarose gel electrophoresis (90 V, 1.5 h).

**Fig. 6.**
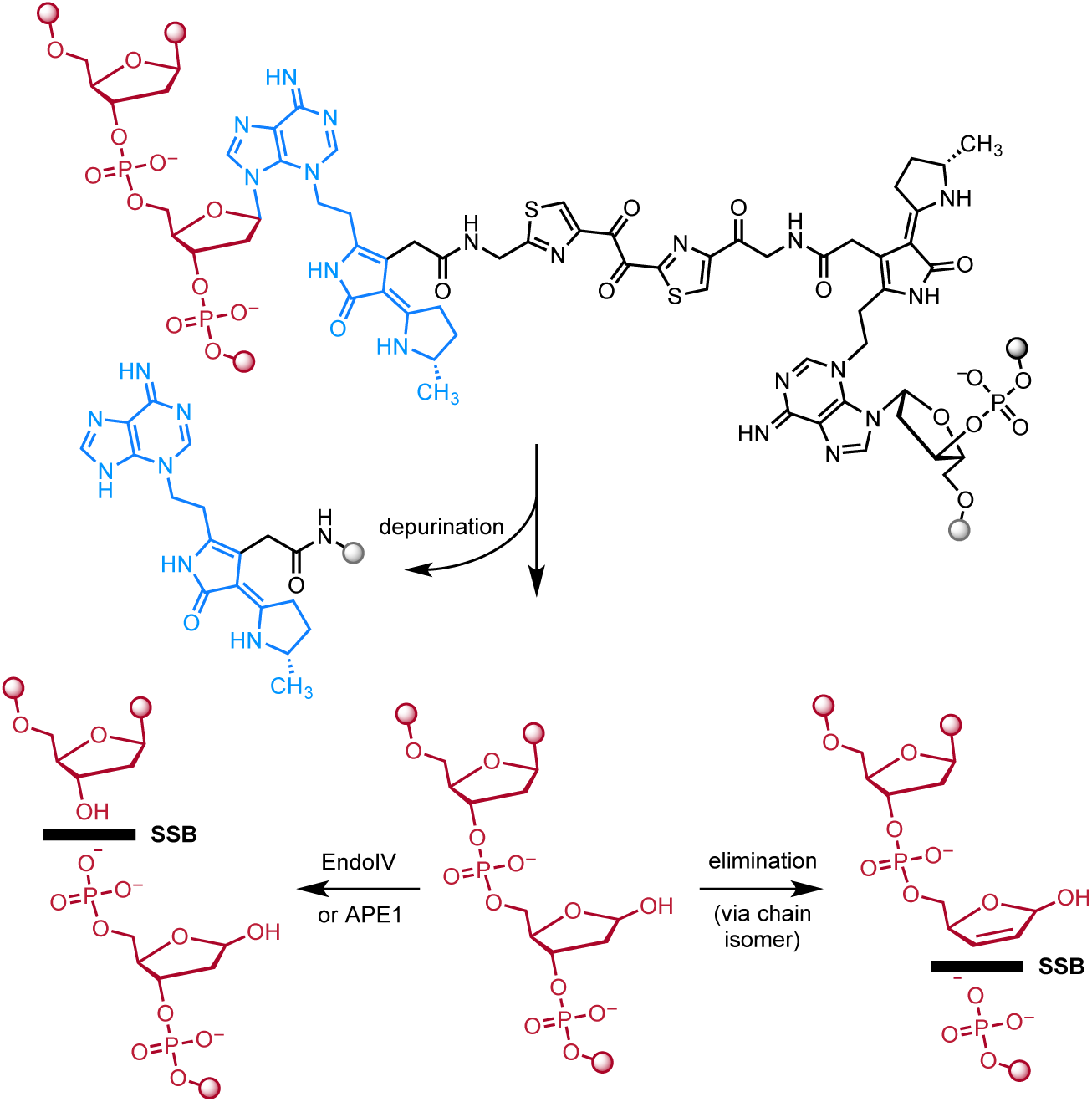
Mechanism of SSB and DSB induction following formation of ICLs. Depurination followed by slower elimination of the 3′ phosphate would introduce a SSB. Alternatively, hydrolysis of the AP site by APE1 in tissue culture would introduce a break in the DNA. Repetition of either pathway at the remaining intact alkylated nucleotide would form a DSB.

## Results

To probe for SSB and DSB formation we began by evaluating the effect of *clb*^+^ BW25113 *E. coli* (*clb*^+^) on circular pUC19 plasmid DNA. Nicking (SSB) of the plasmid results in relaxation of the supercoiled structure and lower mobility on an electrophoretic gel. Cleaved or linearized (DSB) plasmid also has a distinct mobility. *E. coli* lacking the pathway (*clb*^−^) or mutated in the gene *clbL* (S179A) were employed as controls. ClbL is an amidase that has been proposed to mediate a heterodimerization in the final steps of colibactin (**1**) biosynthesis.^5,9^ Thus we anticipated that the DNA ICL phenotype would be abrogated in this strain if colibactin (**1**) were the causative agent. Bacteria were incubated with DNA for 1–9 h and the DNA samples were purified and analyzed by native gel electrophoresis (Fig. 2). A background level of nicking and cleavage was observed in the *clb*^−^ strain. However, this damage was markedly increased in the *clb*^+^ culture after incubating for 4 h (Lane 11), and all of the circular plasmid had converted to nicked (SSB) or linearized (DSB) DNA after 6 h (Lane 12). The DNA damage in the *clbL* mutant (S179A) was comparable to the *clb*^−^ background.

Re-analysis of the samples by denaturing electrophoresis revealed a different DNA migration pattern and a significant amount of degradation of the *clb*^+^ *-*treated DNA after incubating for 4 h (Fig. 3, Lane 11). Complete degradation of the plasmid was observed after 9 h (Lane 13).

The amount of DNA SSBs and DSBs observed from the native gel electrophoresis could be quantified by image analysis. The amount of nicked and cleaved DNA was normalized relative to untreated linearized pUC19 DNA in a control lane (Fig. 4). Data are the average of three technical replicates. A calibration curve correlating the DNA band intensity and the quantity of DNA in the band is shown in Fig. S1. These data show that the number of SSBs significantly increased at 4 h and began to decrease thereafter (Fig. 4A). The number of DSBs increased with progression of time (Fig. 4B). Throughout the time period 0−9 h, smaller amounts of DSBs and SSBs were observed in the co-culture with *clb*^−^ *E. coli* (Figs. 4A, B).

Based on our previous preparations of mass spectrometry samples, 4 h is the optimum incubation time for preparing cross-linked linearized pUC19 DNA using *clb*^+^ *E. coli*.^5,16^ To confirm that these circular pUC19 DNA contained ICLs, we incubated the plasmid with *clb*^+^ *E. coli* for 4 h, and then linearized the DNA using the EcoRI restriction enzyme. The DNA was re-purified and then analyzed using both native and denaturing gel electrophoresis. Fig. 5A shows that linearization of either the *clb*^+^ or *clb*^−^ treated DNA, followed by native gel electrophoresis, produced the expected duplexed DNA (Lanes 7 and 10, compare to Lanes 3 and 4). By comparison, Fig. 5B shows that when the same sample was analyzed by denaturing electrophoresis, only the *clb*^+^-treated DNA remained covalently connected, as evidenced by co-migration with pUC19 DNA that was linearized prior to treatment with *clb*^+^ *E. coli* (compare Lanes 10 and 4). ICLs were not detected in the *clbL* mutant (Fig. S3).

The data shown in Figs. 2–5 can be explained by initial formation of an ICL, depurination, and elimination of the 3′ phosphate to sequentially form SSBs and DSBs (Fig. 6). This models is supported by the rates of depurination and subsequent 3′ phosphate elimination *in vitro*. The reported half-life for depurination of 3-methyladenosine in rat liver DNA is 24 h (pH 7.0, 37 °C).^14^ The reported half-life for elimination of the 3′ phosphate from an AP site in circular phage PM2 DNA is 190 h (pH 7.4, 37 °C).^15,17^ Additionally it is possible that APE1 introduces additional nicks at AP sites in tissue culture. We note that the site of adenine alkylation on the right hand-side of colibactin has not been rigorously established but alkylation at N3 would be consistent with the structure of **3** and the reactivity observed here. N7A adducts are labile toward depurination (t_1/2_ = 3 h), while N6A adducts are stable.^15^

According to this model and the relative rates of depurination and 3′ phosphate elimination, the DNA should be accumulating AP sites. To test this, we exposed the *clb*^+^-treated plasmid DNA to Endonuclease IV (EndoIV) which recognizes AP sites and cleaves the phosphodiester backbone 5′ to the AP site, resulting in a one nucleotide gap flanked by a 3′ hydroxyl and 5′ deoxyribose phosphate.^18^ If AP sites are accumulating, one would expect EndoIV treatment to amplify the amount of DNA breaks. As shown in Fig. 7, all of the plasmid DNA was converted to nicked (SSBs) and linearized (DSBs) DNA after treatment with EndoIV, and further degraded DNA fragments were observable (Lane 9). By comparison, DNA that had been exposed to buffer alone partially converted to nicked (SSBs) DNA, and retained a smaller amount of intact plasmid DNA (Lane 8) comparing to the untreated sample (Lane 7). This is consistent with the mechanism outlined in Fig. 6 where SSBs and DSBs derive from spontaneous depurination, followed by phosphate elimination. The *clb*^−^ and *clbL* mutant (S179A) strains did not display measurable changes in DNA topology on treatment with Endo IV or buffer alone within 20 h of incubation at 37 °C. DNA cleavage increased as the amount of time the *clb*^+^-treated circular or linearized pUC19 DNA was exposed to buffer or EndoIV was increased (Fig S4−S5). However, the circular and linearized untreated pUC19 DNA, or the DNA treated with the 100 µM cisplatin (which forms more stable^15^ N7G ICLs^19^) did not show any further damage over time in the presence or absence of EndoIV (Fig S6−S9). These results indicate that the colibactin-cross-linked DNA undergoes depurination and then evolves more slowly to strand breaks. These breaks are amplified by exposure to Endo IV.

**Fig. 7.**
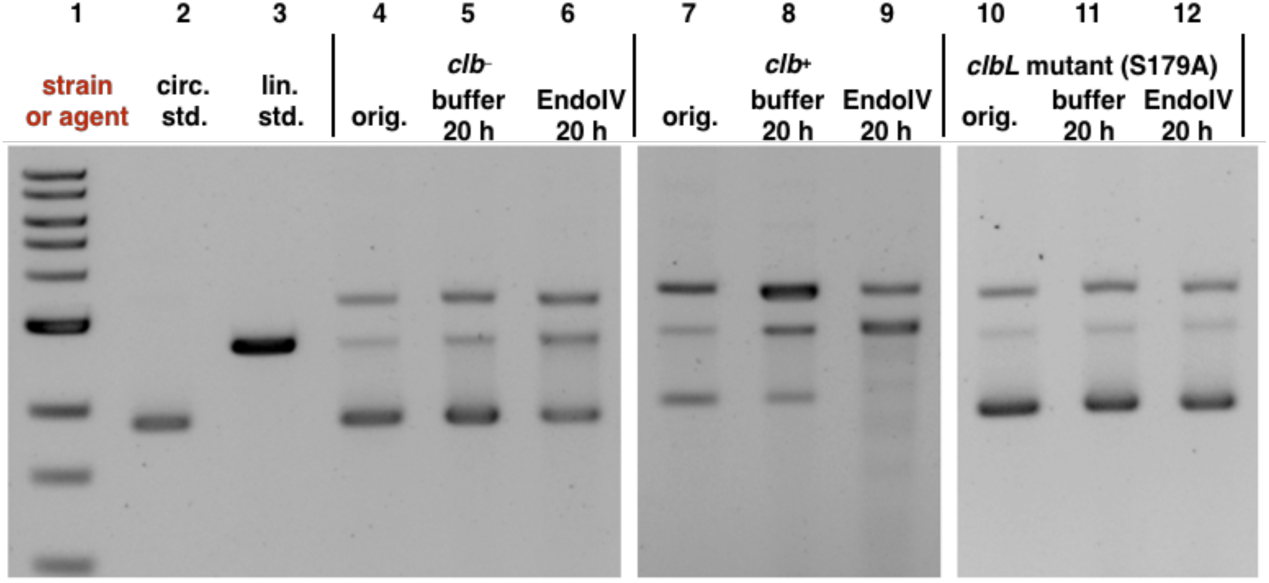
Exposure of pUC19 plasmid DNA to *clb*^+^, followed by incubation with Endo IV leads to consumption of undamaged plasmid and formation of nicked and linearized DNA. This is not observed in the *clb*^−^ or *clbL* mutant controls. DNA ladder (Lane #1); circular pUC19 DNA standard (Lane #2); linearized pUC19 DNA standard (Lane # 3); circular pUC19 DNA isolated from co-culture with *clb*^−^ BW25113 *E. coli* (Lane #4), reacted with buffer (Lane #5), reacted with Endonuclease IV (Lane #6); circular pUC19 DNA isolated from co-culture with *clb*^+^ BW25113 *E. coli* (Lane #7), reacted with buffer (Lane #8), reacted with Endonuclease IV (Lane #9); circular pUC19 DNA isolated from co-culture with *clbL* mutant (S179A) BW25113 *E. coli* (Lane #10), reacted with buffer (Lane #11), reacted with Endonuclease IV (Lane #12). Conditions (Lane #4–#6): circular pUC19 DNA from co-culture with *clb*^−^ BW25113 *E. coli* in M9-CA media for 4 h at 37 °C (Lane # 4); the DNA (3.9 µM base pair) was reacted with NEBuffer 3.1 (New England Biolabs®), pH 7.9, at 37 °C for 20 hours (Lane #5); the DNA (3.9 µM base pair) was further reacted with 20 units of Endonuclease IV in NEBuffer 3.1® (New England Biolabs®), pH 7.9, at 37 °C for 20 hours (Lane #6). Conditions (Lane #7–#9): circular pUC19 DNA isolated from co-culture with *clb*^+^ BW25113 *E. coli.* in in M9-CA media for 4 h at 37 °C (Lane #7); the DNA (3.9 µM base pair) was reacted with NEBuffer 3.1 (New England Biolabs®), pH 7.9, at 37 °C for 20 hours (Lane #8); the DNA (3.9 µM base pair) was reacted with 20 units of Endonuclease IV in NEBuffer 3.1® (New England Biolabs®), pH 7.9, at 37 °C for 20 hours (Lane #9). Conditions (Lane #10–#12): circular pUC19 DNA isolated from co-culture with *clbL* mutant (S179A) BW25113 *E. coli.* in M9-CA media for 4 h at 37 °C (Lane #10); the DNA (3.9 µM base pair) was reacted with NEBuffer 3.1® (New England Biolabs®), pH 7.9, at 37 °C for 20 hours (Lane #11); the DNA (3.9 µM base pair) was reacted with 20 units of Endonuclease IV in NEBuffer 3.1® (New England Biolabs®), pH 7.9, at 37 °C for 20 hours (Lane #12). The DNA was not re-purified and was directly analyzed by native agarose gel electrophoresis (90 V, 1.5 hr).

We then sought to determine if these effects were recapitulated by synthetic material. Because the carbon–carbon bond within the α-diketone function of colibactin (**1**) is susceptible to nucleophilic cleavage^20^ we employed the enediol **4** in our studies (Fig. 8). We have shown that oxidation to the α-dicarbonyl occurs spontaneously under air,^20^ and that cyclodehydration in aqueous buffer rapidly forms the DNA warheads of **1**.^6^ We evaluated the effects of **4** on DNA. The DNA was completely degraded at 100 µM **4** (Fig. 9A Lane 5; Fig. 9B, Lane 3). Fig. 9A shows that ICLs are formed when linearized pUC19 DNA is exposed to 10 µM–100 nM of **4** for 4 h (compare Lanes 6–8 to Lane 4). Lanes 3–5 of Fig. 9B show that most of the circular pUC19 DNA remains intact following exposure to 10 or 1 µM **4** for 3 h and analysis by native gel electrophoresis. However, on treatment with EndoIV, significant amounts of nicked and linearized DNA are produced (Lanes 8 and 9). The degradation of DNA was qualitatively higher than that obtained when incubating in buffer alone (Lanes 6 and 7). These studies show that synthetic **4** recapitulates the activity of *clb*^+^ *E. coli* in this assay.

**Fig. 8.**
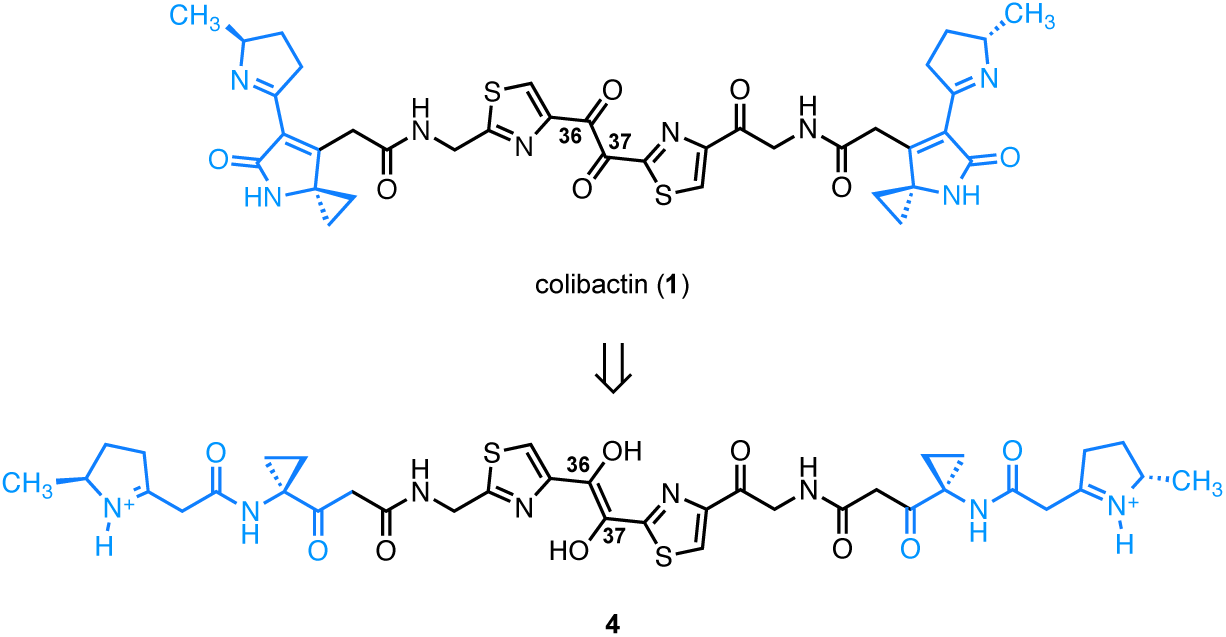
Structure of the colibactin precursor **4**.

**Fig. 9.**
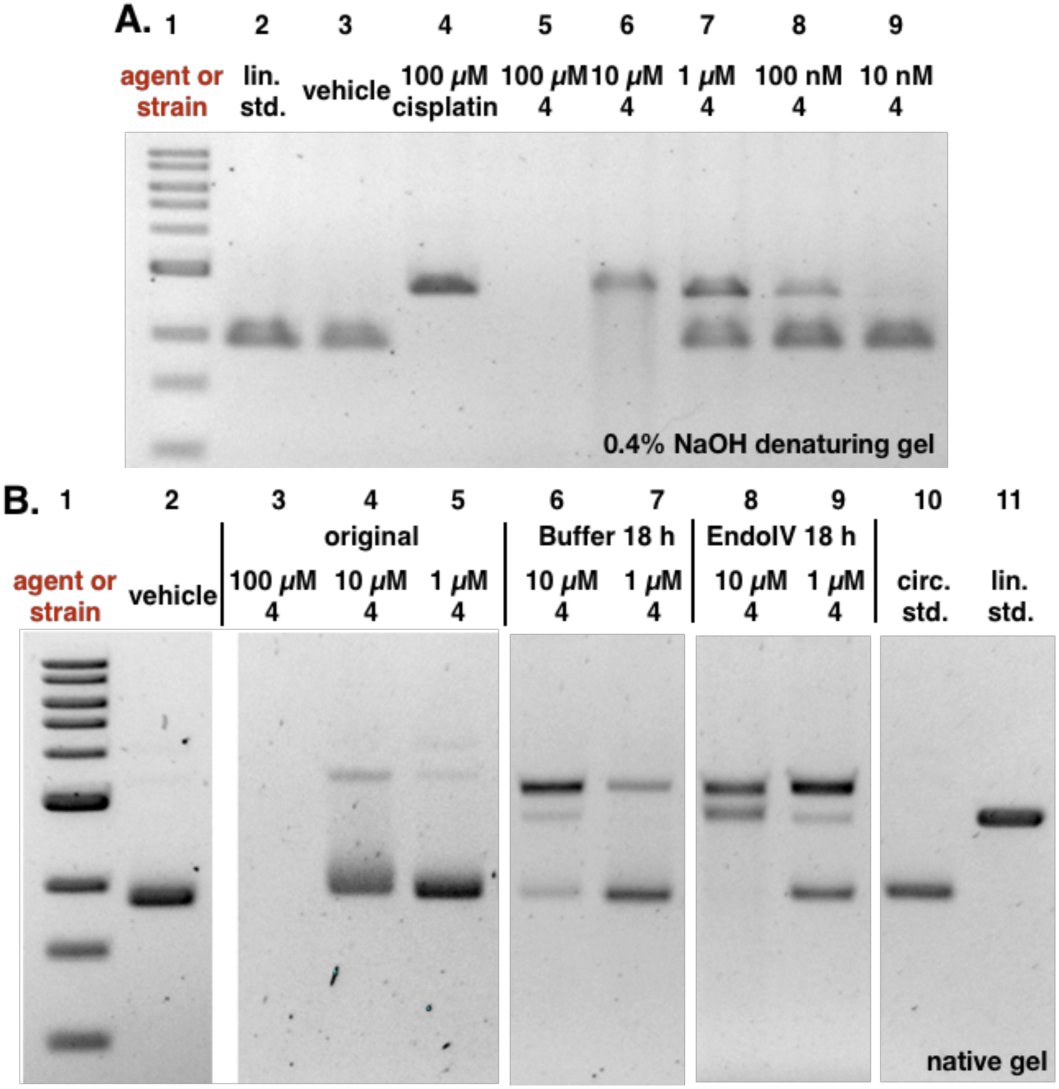
Analysis of induction of AP sites by the colibactin precursor **4. A.** Incubation of plasmid pUC19 DNA exposed to **4** in buffer for 18 h results in minor nicking and cleavage. **B.** Addition of EndoIV increases the amount of nicked and cleaved plasmid. Conditions: A. 5% DMSO was used as vehicle (negative control), and 100 µM cisplatin was used as positive control. DNA ladder (Lane #1); linearized pUC19 DNA standard (Lane #2); 5% DMSO (Lane #3); 100 µM cisplatin (Lane #4); 100 µM **4** (Lane #5); 10 µM **4** (Lane #6); 1 µM **4** (Lane #7); 100 nM **4** (Lane #8); 10 nM **4** (Lane #9). Conditions (Lane #3): linearized pUC19 DNA (15.4 µM in base pairs), 5% DMSO (vehicle), 10 mM citric buffer, pH 5.0, 4 h, 37 °C. Conditions (Lane #4): linearized pUC19 DNA (15.4 µM in base pairs), 5% DMSO (vehicle), 100 µM cisplatin, 10 mM citric buffer, pH 5.0, 4 h, 37 °C. Conditions (Lanes #5–#9): circular pUC19 DNA (15.4 µM in base pairs), **4** (100 µM–10 nM), 5% DMSO, 10 mM citric buffer, pH 5.0, 4 h, 37 °C. The DNA was analyzed by 0.4% NaOH denaturing agarose gel electrophoresis (90 V, 1.5 h). B. 5% DMSO was used as vehicle. DNA ladder (Lane #1); 5% DMSO (Lane #2); 100 µM **4** (Lane #3); 10 µM **4** (Lane #4); 1 µM **4** (Lane #5); post buffer-reacted after 10 µM **4** (Lane #6); post buffer-reacted after 1 µM **4** (Lane #7); post EndoIV-reacted after 10 µM **4** (Lane #8); post EndoIV-reacted after 1 µM **4** (Lane #9); circular pUC19 plasmid standard (Lane #10); linearized pUC19 plasmid standard (Lane #11). Conditions (Lane #2): circular pUC19 DNA (15.4 µM in base pairs), 5% DMSO (vehicle), 10 mM citric buffer, pH 5.0, 4 h, 37 °C. Conditions (Lanes #3–#5): circular pUC19 DNA (15.4 µM in base pairs), **4** (100 µM–1 µM), 5% DMSO, 10 mM citric buffer, pH 5.0, 3 h, 37 °C. Conditions (Lanes #6–#7): **4** (10 µM–1 µM)-treated circular pUC19 DNA (3.9 µM in base pairs), NEBuffer 3.1® (New England Biolabs®), pH 7.9, at 37 °C for 18 h. Conditions (Lanes #8–#9): **4** (10 µM–1 µM)-treated circular pUC19 DNA (3.9 µM in base pairs), 20 units of Endonuclease IV (New England Biolabs®), NEBuffer 3.1® (New England Biolabs®), pH 7.9, at 37 °C for 18 h. The DNA was analyzed by native agarose gel electrophoresis (90 V, 2 h).

## Discussion

This data provides molecular and mechanistic knowledge of colibactin induced DNA damage. We show that colibactin derived DNA crosslinks are inherently unstable and provide a molecular model that explains the evolution of colibactin ICLs to DSBs. Our data explain how colibactin can induce two discreet biological phenotypes simultaneously: (1) DSBs via ICL repair and (2) DSBs formed independently of the ICL repair pathway.

A prior study established that *clb*^+^ *E. coli* form ICLs and activate the FA ICL repair pathway.^8^ FA repair, in turn, leads to the generation of DNA DSBs. FA-dependent DSBs are resolved by the HR DNA DSB repair pathway.^10^ Thus the generation of ICLs is consistent with the earlier determination that *clb*^+^ *E. coli* lead to accumulation of the HR repair factor γH2AX in eukaryotic cells, fragmentation of genomic DNA in a neutral comet assay,^7^ and co-localization of the FA repair protein Fanconi anemia protein D2 (FANCD2) with γH2AX.^8^

However cells deficient in Ku80, a factor in the non-homologous end joining (NHEJ) DSB repair pathway,^11^ are sensitized to *clb*^+^ *E. coli.*^21^ Because NHEJ is not coupled to FA, this suggests a potential additional mechanism for DSB generation. We had observed that DNA alkylation products derived from synthetic colibactin analogs and colibactin (**1**) itself were unstable.^5-6,16b^ We hypothesized that this inherent ICL instability could explain the sensitization of Ku80 deficient cells to colibactin genotoxicity. To probe this in more detail, we studied the stability of alkylation products derived from circular pUC19 DNA. Here we used the topology of the supercoiled plasmid DNA as a proxy for the stability of colibactin alkylation products. Depurination of a single alkylation product followed by elimination of the 3′ phosphate leads to a SSB (nick) and relaxation of the supercoiled plasmid. Repetition of the same sequence on the complementary strand leads to linearization of the plasmid.

Our data suggest that the colibactin-derived ICLs are unstable and undergo spontaneous depurination and subsequent elimination of the 3′ phosphate. The amount of nicking and cleavage of DNA was increased on treatment with EndoIV, suggesting the accumulation of AP sites, followed by slower strand cleavage. This is consistent with the relative rates of depurination and 3′ phosphate elimination (t_1/2_ = 24 and 190 h respectively, vide supra). Because the rate of 3′ phosphate elimination is slow, we speculate that the recognition of AP sites by endonucleases, such as APE1, may be contribute to DNA break formation in tissue culture.

This work provides several other insights into colibactin-mediated DNA damage. First, as shown in Fig 7, EndoIV treatment did not increase the number of SSBs and DSBs in DNA exposed to the *clbL* mutant (S179A). We have previously shown that a *clbL* mutant (S179A) alkylates but does not cross-link DNA.^22^ This suggests that the alkylated bases within the colibactin ICL are less stable than those derived from monoalkylation products. This may derive from distortion of the DNA duplex, as observed with other ICL agents such as cisplatin.^19^ Additionally while the site of DNA alkylation by colibactin (**1**) has not been rigorously established, the monoalkylation product **3** is linked through N3A.^13^ Both N3 and N7 adenine adducts are unstable,^15^ suggesting ICLs are formed by alkylation at N3,N3 or N3,N7.

## Material and Methods

### Bacterial Strains and Growth Conditions

The *E. coli* strains used in this study were prepared by the Crawford laboratory at Yale University (New Haven, CT). The BW25113 *E. coli.* contain either the colibactin gene cluster on pBeloBac11 (*clb*^+^), the empty vector alone (*clb*^−^), or the vector carrying the *clbL* point mutant (S179A). The *clbL* point mutant (S179A) was constructed as previously described.^23^ Bacteria were grown overnight in LB media containing chloramphenicol (25 µg/mL), and then the grown cultures were diluted into M9-cas-amino acid (M9-CA) medium. To prepare the M9-CA medium, the M9 minimum media (Difco) was supplemented with 0.4% glucose, 2 mM MgSO_4_, 0.1 mM CaCl_2_, 12.5 µg/mL chloramphenicol and the following L-amino acid mass composition (5 g/L total): 3.6% Arg, 21.1% Glu, 2.7% His, 5.6% Ile, 8.4% Leu, 7.5% Lys, 4.6% Phe, 9.9% Pro, 4.2% Thr, 1.1% Trp, 6.1% Tyr, 5% Val, 4% Asn, 4% Ala, 4% Met, 4% Gly, 4% Cys, and 4% Ser. Standard growth conditions were 37 °C with 250 rpm agitation.

### Synthetic and Commercial Reagents

Cisplatin was purchased from Biovision® (Milpitas, CA). The synthetic colibactin linear precursor (**4**) was prepared by a modification of the published procedure.^5^

### In vitro Plasmid Cleavage Assays

The 2686 bp pUC19 vector (circular) was purchased from New England Biolabs® (Ipswich, MA). For each reaction with *E.coli*, 1000 ng of circular pUC19 DNA was added to 200 µL (7.7 µM base pairs) of M9-CA medium inoculated with 1.2 × 10^7^ bacteria pre-grown to exponential phase in the M9-CA medium. The mixture of DNA and bacteria was incubated at 37 °C for 1 h−9 h, and the bacteria were then pelleted. The DNA was isolated from the supernatant using the polymerase chain reaction (PCR) clean-up kit (New England Biolabs®) and quantified using a nanodrop. The DNA was then stored in −20 °C until electrophoretic analysis or follow-up experiments.

For each reaction with synthetic colibactin linear precursor (**4**), 200 ng of circular pUC19 DNA (15.4 µM base pairs) was incubated with **4** in concentrations from 100 µM−1 nM, in a total volume of 20 µL. **4** was diluted in DMSO such that each reaction contained 5% DMSO concentration. Reactions were conducted in 10 mM sodium citrate buffer (pH 5.0). Pure DMSO vehicle was used as negative control. A control employing 200 ng of circular pUC19 DNA (15.4 µM base pairs) was treated with pure DMSO vehicle in 10 mM sodium citrate (pH 5) buffer with a final DMSO concentration of 5% was prepared simultaneously. Reactions proceeded for 4 h at 37 °C, and the reacted DNA was stored at −20 °C until electrophoretic analysis or follow-up experiments.

### DNA Cross-linking Assays

Linearized pUC19 DNA was used for all DNA cross-linking assays. To prepare the linearized DNA, the 2686 bp pUC19 vector (New England Biolabs®) was linearized with 20 units/µg EcoRI-HF® (New England Biolabs®) and the linearized DNA was purified using a PCR clean-up kit (New England Biolabs®), eluted into 10 mM Tris (pH 8.0), and quantified using a nanodrop. For each reaction with *E.coli*, 800 ng of linearized pUC19 DNA was added to 200 µL (6.2 µM base pairs) of M9-CA medium inoculated with 1.2 × 10^7^ bacteria pre-grown to exponential phase in the M9-CA medium. The mixture of DNA and bacteria was incubated at 37 °C for 4 h, and the bacteria were then pelleted. The DNA was isolated from the supernatant using the PCR clean-up kit (New England Biolabs®) and quantified using a nanodrop. The DNA was then stored at −20 °C until electrophoretic analysis or follow-up experiments.

For each reaction with synthetic colibacin linear precursor (**4**), 200 ng of linearized pUC19 DNA (15.4 µM base pairs) was incubated with **4** in concentrations ranging from 100 µM−1 nM in a total volume of 20 µL. **4** was diluted in DMSO such that each reaction contained a fixed 5% DMSO concentration. Reactions were conducted in 10 mM sodium citrate buffer (pH 5.0). Pure cisplatin (Biovision®) and DMSO vehicle were used as positive and negative controls, respectively. Stock solutions of cisplatin in DMSO were prepared immediately before use. A control containing 200 ng of circular pUC19 DNA (15.4 µM base pairs) was treated with 100 µM cisplatin or DMSO vehicle in 10 mM sodium citrate (pH 5) buffer with a final DMSO concentration of 5% was prepared simultaneously. Reactions were conducted for 4 h at 37 °C, The DNA was stored at −20 °C until electrophoretic analysis or follow-up experiments.

### Plasmid Linearization Test

The pUC19 DNA isolated from plasmid cleavage assays with *E. coli.* was used for the plasmid linearization test. To set up the linearization reactions, 20 units of EcoRI-HF® (New England Biolabs®) was mixed with 500 ng of isolated DNA (40 units/µg DNA) in CutSmart® buffer (New England Biolabs®), pH 7.9, in a total volume of 50 µL for 30 min at 37 °C. The CutSmart® buffer (New England Biolabs®) contains 50 mM potassium acetate, 20 mM Tris-acetate, 10 mM magnesium acetate, and 100 µg/mL BSA. To set up the negative control reactions, 500 ng of isolated DNA was treated with CutSmart buffer® (New England Biolabs®), pH 7.9, in a total volume of 50 µL for 30 min at 37 °C. The reacted DNA was then purified using PCR clean-up kit (New England Biolabs®) and quantified using the nanodrop. The DNA was then stored at –20 °C before electrophoretic analysis. The *clb*^+^ cross-linked linearized pUC19 DNA was used as positive control for analysis.

### Endonuclease IV Stability Test

The pUC19 DNA isolated from plasmid cleavage and DNA cross-linking assays with *E. coli.* was used for the Endonuclease IV (EndoIV) stability tests. The pUC19 DNA exposed to synthetic compounds in plasmid cleavage or DNA cross-linking assays was directly diluted and used for EndoIV stability tests. To set up each reaction, 50 ng of processed DNA was mixed with 20 units of EndoIV in NEBuffer 3.1® (New England Biolabs®), pH 7.9, in a total volume of 20 µL for 16 h−20 h (unless otherwise noted) at 37 °C. The NEBuffer 3.1® (New England Biolabs®) contained 100 mM sodium chloride, 50 mM Tris-HCl, 10 mM magnesium chloride, and 100 µg/ml BSA. To set up each negative control, 50 ng of processed DNA was mixed with NEBuffer 3.1® (New England Biolabs®), pH 7.9, in a total volume of 20 µL for 16 h−20 h (otherwise otherwise noted) at 37 °C. Following completion of the experiment, the DNA was stored at –20 °C before electrophoretic analysis.

### Gel Electrophoresis

For each DNA sample (except the samples employed in the EndoIV stability tests), the DNA concentration was pre-adjusted to 10 ng/µL. Five microliters (50 ng) of DNA was removed and mixed with 1.5 µL of 6 × purple gel loading dye; no SDS (New England Biolabs®), and loaded onto 1% agarose Tris Borate EDTA (TBE) gels. For the samples employed in the EndoIV stability tests, the DNA solution (20 µL) was mixed with 4 µL of 6 × purple gel loading dye; no SDS (New England Biolabs®) and directly loaded onto 1% agarose Tris Borate EDTA (TBE) gels. For denaturing gels, 5 µL (50 ng) of DNA was removed each time and separately mixed with 15 µL of 0.2% denaturing buffer (0.27% sodium hydroxide, 10% glycerol, and 0.013% bromophenol blue) or 0.4% denaturing buffer (0.53% sodium hydroxide, 10% glycerol, and 0.013% bromophenol blue) in an ice bath. The mixed DNA samples were denatured at 4 °C for 10 min and then immediately loaded onto 1% agarose Tris Borate EDTA (TBE) gels. All gel electrophoresis was conducted at 90 V for 2 h (unless otherwise noted). The gel was stained with SybrGold (Thermo Fisher) for 2 h.

### Quantification of Gel Images

The gel images were quantified using Adobe®Photoshop® CC 2018. The pixel intensity of each gel band was carefully measured, and the average background pixel intensity was deducted from each gel band’s raw measurement. The net pixel intensity values of each target gel band were normalized by the net pixel intensity of the linearized pUC19 standard. A standard curve of the DNA quantity vs. net pixel intensity is included in the Supporting Information.

## Acknowledgement

Financial support from the National Institutes of Health (R01CA215553) is gratefully acknowledged. We thank Emilee Shine (Yale University) for providing the *clbL* mutant.

## Author contributions

M.Z. designed and performed the DNA alkylation experiments. K.M.W. prepared **4**. S.B.H. and M.Z. wrote the manuscript, K.M.W. provided comments on the manuscript.

## References

1. (a)Balskus, E. P. Nat. Prod. Rep. 2015, 32, 1534. (b)Trautman, E. P.; Crawford, J. M. Curr. Top. Med. Chem. 2015, 16, 1. (c)Taieb, F.; Petit, C.; Nougayrede, J. P.; Oswald, E. EcoSal Plus 2016, 7, doi: 10.1128/ecosalplus.ESP. (d)Healy, A. R.; Herzon, S. B. J. Am. Chem. Soc. 2017, 139, 14817. (e)Faïs, T.; Delmas, J.; Barnich, N.; Bonnet, R.; Dalmasso, G. Toxins 2018, 10, 151.

2. Putze, J.; Hennequin, C.; Nougayrede, J. P.; Zhang, W.; Homburg, S.; Karch, H.; Bringer, M. A.; Fayolle, C.; Carniel, E.; Rabsch, W.; Oelschlaeger, T. A.; Oswald, E.; Forestier, C.; Hacker, J.; Dobrindt, U. Infect. Immun. 2009, 77, 4696.

3. Arthur, J. C.; Perez-Chanona, E.; Mühlbauer, M.; Tomkovich, S.; Uronis, J. M.; Fan, T.-J.; Campbell, B. J.; Abujamel, T.; Dogan, B.; Rogers, A. B.; Rhodes, J. M.; Stintzi, A.; Simpson, K. W.; Hansen, J. J.; Keku, T. O.; Fodor, A. A.; Jobin, C. Science 2012, 338, 120.

4. Buc, E.; Dubois, D.; Sauvanet, P.; Raisch, J.; Delmas, J.; Darfeuille-Michaud, A.; Pezet, D.; Bonnet, R. PLoS One 2013, 8, e56964.

5. Xue, M.; Kim, C. S.; Healy, A. R.; Wernke, K. M.; Wang, Z.; Frischling, M. C.; Shine, E. E.; Wang, W.; Herzon, S. B.; Crawford, J. M. Science 2019, 365, eaax2685.

6. Healy, A. R.; Nikolayevskiy, H.; Patel, J. R.; Crawford, J. M.; Herzon, S. B. J. Am. Chem. Soc. 2016, 138, 15563.

7. Nougayrède, J.-P.; Homburg, S.; Taieb, F.; Boury, M.; Brzuszkiewicz, E.; Gottschalk, G.; Buchrieser, C.; Hacker, J.; Dobrindt, U.; Oswald, E. Science 2006, 313, 848.

8. Bossuet-Greif, N.; Vignard, J.; Taieb, F.; Mirey, G.; Dubois, D.; Petit, C.; Oswald, E.; Nougayrede, J. P. Mbio 2018, 9, e02393.

9. (a)Jiang, Y.; Stornetta, A.; Villalta, P. W.; Wilson, M. R.; Boudreau, P. D.; Zha, L.; Balbo, S.; Balskus, E. P. bioRxiv 2019, 567248. (b)Jiang, Y.; Stornetta, A.; Villalta, P. W.; Wilson, M. R.; Boudreau, P. D.; Zha, L.; Balbo, S.; Balskus, E. P. J. Am. Chem. Soc. 2019, 141, 11489.

10. Clauson, C.; Schärer, O. D.; Niedernhofer, L. Cold Spring Harbor Perspect. Biol. 2013, 5,

11. Scully, R.; Panday, A.; Elango, R.; Willis, N. A. Nat. Rev. Mol. Cell Biol. 2019, 20, 698.

12. Michl, J.; Zimmer, J.; Tarsounas, M. The EMBO Journal 2016, 35, 909.

13. Wilson, M. R.; Jiang, Y.; Villalta, P. W.; Stornetta, A.; Boudreau, P. D.; Carra, A.; Brennan, C. A.; Chun, E.; Ngo, L.; Samson, L. D.; Engelward, B. P.; Garrett, W. S.; Balbo, S.; Balskus, E. P. Science 2019, 363, eaar7785.

14. (a) Margison, G. P.; O’Connor, P. J. Biochim. Biophys. Acta, Nucleic Acids Protein Synth. 1973, 331, 349. (b)Fujii, T.; Saito, T.; Nakasaka, T. Chem. Pharm. Bull. 1989, 37, 2601.

15. Gates, K. S. Chem. Res. Toxicol. 2009, 22, 1747.

16. (a)Xue, M.; Shine, E.; Wang, W.; Crawford, J. M.; Herzon, S. B. Biochemistry 2018, 57, 6391. (b)Xue, M.; Kim, C. S.; Healy, A. R.; Wernke, K. M.; Wang, Z.; Frischling, M. C.; Shine, E. E.; Wang, W.; Herzon, S. B.; Crawford, J. M. bioRxiv 2019, 574053.

17. (a)Lindahl, T.; Andersson, A. Biochemistry 1972, 11, 3618. (b)Crine, P.; Verly, W. G. Biochim Biophys Acta 1976, 442, 50.

18. (a)Ljungquist, S. J. Biol. Chem. 1977, 252, 2808. (b)Levin, J. D.; Johnson, A. W.; Demple, B. J. Biol. Chem. 1988, 263, 8066.

19. Coste, F.; Malinge, J. M.; Serre, L.; Shepard, W.; Roth, M.; Leng, M.; Zelwer, C. Nucleic Acids Res. 1999, 27, 1837.

20. Healy, A. R.; Wernke, K. M.; Kim, C. S.; Lees, N. R.; Crawford, J. M.; Herzon, S. B. Nat. Chem. 2019, 11, 890.

21. Cuevas-Ramos, G.; Petit, C. R.; Marcq, I.; Boury, M.; Oswald, E.; Nougayrède, J.-P. Proc. Natl. Acad. Sci. U. S. A. 2010, 107, 11537.

22. Shine, E. E.; Xue, M.; Patel, J. R.; Healy, A. R.; Surovtseva, Y. V.; Herzon, S. B.; Crawford, J. M. ACS Chem. Biol. 2018, 13, 3286.

23. Trautman, E. P.; Healy, A. R.; Shine, E. E.; Herzon, S. B.; Crawford, J. M. J. Am. Chem. Soc. 2017, 139, 4195.

